# The swim-up technique separates bovine sperm by metabolic rates, motility and tail length

**DOI:** 10.1101/624502

**Authors:** Veronika Magdanz, Sergii Boryshpolets, Clara Ridzewski, Barbara Eckel, Klaus Reinhardt

**Affiliations:** Chair of Applied Zoology, TU Dresden Zellescher Weg 20, 01217 Dresden, Germany; University of South Bohemia in České Budějovice Faculty of Fisheries and Protection of Waters, South Bohemian Research Center of Aquaculture and Biodiversity of Hydrosensors Zátiší 728/II, 389 25 Vodňany, Czech Republic

**Keywords:** swim-up, bovine sperm, metabolism, sperm selection, sperm quality

## Abstract

Swim-up is a sperm purification method that is being used daily in andrology labs around the world as a simple step for *in vitro* sperm selection. This method accumulates the most motile sperm in the upper fraction and leaves sperm with low or no motility in the lower fraction but the underlying reasons are not fully understood. In this article, we compare metabolic rate, motility and sperm tail length of bovine sperm cells of the upper and lower fraction. The metabolic assay platform reveals oxygen consumption rates and extracellular acidification rates simultaneously and thereby delivers the metabolic rates in real time. Our study confirms the upper fraction of bull sperm has improved motility compared to the cells in the lower fraction and shows higher metabolic rates. This pattern was consistent across media of two different levels of viscosity. Sperm with longer flagella are selected in the upper fraction. We conclude that the motility-based separation of the swim-up technique is based on metabolic differences. Metabolic assays could serve as additional or alternative, label-free method to evaluate sperm quality, which is likely particularly useful in cases of asthenozoospermia and teratospermia. Furthermore, metabolic measurements of sperm cells can reveal differences in metabolic pathways in different environments.

## Introduction

Infertility affects 10-15% of couples worldwide, and about 15 % of these cases remain unexplained^1,2^. Among various other processes, successful fertilization requires the migration of sperm through the female reproductive tract, a process hard to reconstruct *in vitro*. Except for sperm morphology, current i*n vitro* sperm selection assays in assisted reproduction are mostly based on motility assays, maturity (based on sperm binding to hyaluronic acid), zona pellucida binding, assessment of acrosome reaction, or mucus penetration^3,4^ - all processes that require energy.

Sperm cells obtain their energy through two main pathways, oxidative phosphorylation (ATP production through the electron transport chain of the mitochondria) or glycolysis (ATP production in the cytosol by breakdown of sugars). The degree to which these metabolic pathways are employed in sperm is species-specific and either oxidative phosphorylation or glycolysis can be more pronounced under certain conditions^5^. For instance, it is known that in murine sperm, glycolysis is the dominant pathway, while in bovine sperm, oxidative phosphorylation is preferred^5–8^. In human sperm, both pathways are essential for energy production. However, glycolysis is sufficient to maintain motility if glucose is present in the medium^9,10^. It is also predicted that sperm can switch between pathways depending on the conditions in the female reproductive tract, substrate and oxygen concentrations^8,11^.

As most of the sperm’s energy goes towards motility, it is commonly used as an indicator for predicting fertilization success^8,12,13^. In birds and external fertilizers, where higher sperm motility is beneficial under sperm competition, selection for higher energetic capacity in sperm was suggested^14,15^.

Swim-up is a simple, clinical routine method to separate motile from non-motile spermatozoa *in vitro*^16^ whereby sperm are allowed to swim upwards through an overlaid medium. Sperm that do not migrate into the upper medium form the lower fraction. Compared to these sperm, sperm that swim up into the upper medium (upper fraction) displayed increased motility, higher average velocity, higher percentage of normal morphology and generated improved fertilization rates in vitro^17–22^. Therefore, somehow, the swim-up method captures the sperm fraction that also masters the complexity of sperm migration and transport through the reproductive tract to the fertilization site^21^. It is clear that these sperm should be the ones that are enriched to be used in artificial reproduction technologies in humans. However, the underlying mechanism seems yet unknown. This has several important consequences. First, it is not clear to what extent positively selected swim-up sperm with proven increased fertilisation rates in vitro will represent successful sperm migration in the female reproductive tract in vivo. For instance, the swim-up medium is less viscous than the female reproductive tract mucus. During their migration through the reproductive tract, sperm cells encounter a wide range of viscosities, and high viscosity is thought to act as selective agent/barrier by the female. It is also known that the flagellar beating pattern of sperm is altered in high viscosities.^23^ We take this as an incentive to investigate the metabolism of the different sperm swim-up fractions in a medium with viscosity equivalent to the female reproductive tract mucus^24,25^. Second, currently, little seems known whether motility, or swim-up, differences are based on differences in metabolic rate or metabolic pathways. This seems important because under oxidative, but not glycolytic sperm metabolism, sperm damage by oxygen radicals can be expected^26^. This notion may be more important in non-human vertebrates because females of non-human vertebrates can store sperm for days to months, even years^27^ and therefore it is not clear how the short-term success identified by swim-up will translate into fertilisation success of sperm that have been stored. For example, if swim-up success would be caused by a high rate of oxidative sperm metabolism, it is conceivable that this high short-term oxidative expenditure may translate into oxidative damage in the mid- or long-term.

Thus, both, clinical analysis as well as sperm selection in assisted reproduction technologies in humans and animals would benefit from information about the underlying metabolic pathway of the swim-up fractions. Here we compare the metabolic activity of sperm in swim-up fractions in 12 bulls from three genetically different breeds. We perform metabolic measurements before and after swim-up procedure and compare metabolic rates from oxidative phosphorylation and glycolysis of the upper and lower fractions and analyse whether sperm metabolism is repeatable across two levels of viscosity and so predicts motility through environments of different viscosity.

## Results

### Metabolic rates of bovine sperm after swim-up

The rate of oxidative phosphorylation and glycolytic rate are obtained by simultaneously measuring the oxygen consumption rate (OCR) and extracellular acidification rate (ECAR) in an extracellular flux analyzer. The oxygen consumption rate is a direct measure of the mitochondrial electron transport rate and thereby stands as an equivalent for oxidative phosphorylation, during which oxygen is consumed by the cells. The extracellular acidification derives from the lactic acid formed during glycolysis. For a previous application in sperm see Tourmente et al.^6^. The basal rates for OCR and ECAR of the upper, lower and before swim-up fractions were measured for 22 minutes comprising four measurement points each consisting of 15 data points (see Figure 1).

The upper fraction showed higher OCR across the three bulls from three different breeds (Holstein, Fleckvieh, Angus) (Figure 1, yellow data points). The average OCR of upper fraction sperm was up to an order of magnitude higher than in the “before swim-up” and lower fraction sperm population (Figure 1).

Generally, the low ECAR values of all sperm fractions indicate that bull sperm perform little glycolysis. The three bulls differed in the temporal change in glycolytic activity (Figure 1, lower panel). The upper fraction showed higher ECAR activity compared to the “before swim-up” sperm except in the Holstein individual. The basal ECAR was consistently higher in the upper fractions compared to the lower swim-up fractions (blue data sets, Figure 1). Additionally, the ratio between upper and lower fraction OCR and ECAR is different in different individuals.

**Figure 1:**
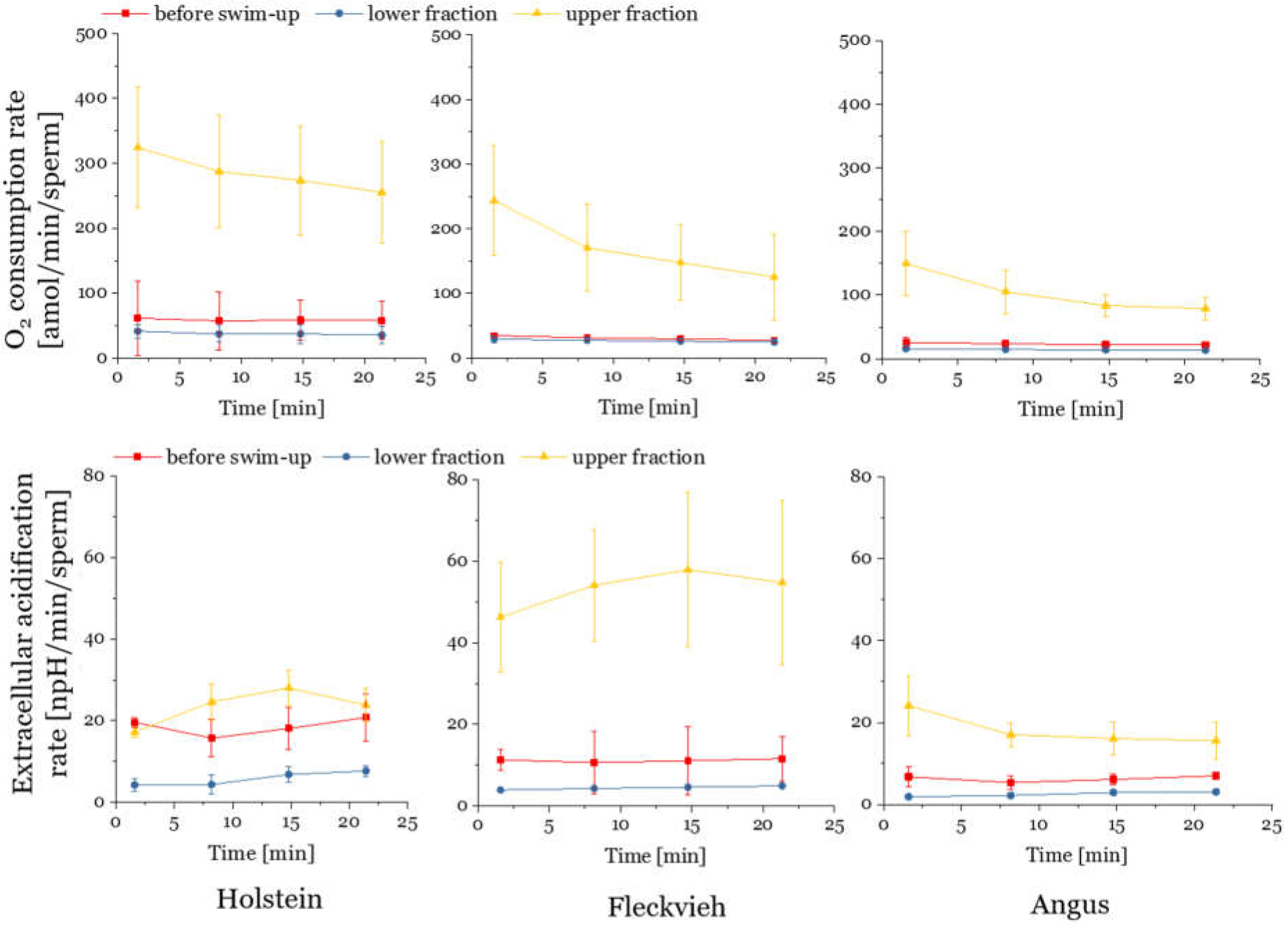
Basal sperm metabolic rates over time in three representative bulls belonging to different breeds. Oxygen consumption rates (OCR) (top) and extracellular acidification rates (bottom) of bull sperm that did not undergo swim-up (before swim-up), upper and lower fractions after swim-up procedure of three bulls of different cattle breeds. Each data point is obtained from a mean value from 3 wells and the data were normalized to the number of cells in each well.

This indicates that the swim-up selects sperm with higher metabolic rates. The fact that the upper fractions display higher OCR and ECAR was not caused by larger sperm numbers in the lower fraction, because all OCR and ECAR values were normalized to the sperm number per well. The density of sperm does not influence their metabolic activity.^28^ On average, the before swim-up fraction contained 4×10^6^ cells/mL, the lower fraction 2×10^7^ cells /mL and the upper fraction 8×10^5^ cells/mL, which means that 23% of cells from the initial sample (before swim-up) migrate to the upper fraction during the swim-up. A low proportion of motile sperm remaining in the lower fraction may cause the OCR in the lower fraction to show lower average values. We tested for this idea in sperm samples from three bulls by normalizing OCR and ECAR to the number of motile sperm in each fraction (Figure S2) even though such normalization has the caveat that metabolically active but non-motile sperm are excluded.

Motility was measured immediately before the metabolic measurements (see Methods). The swim-up accumulates the motile sperm in the upper fraction (motility values of 45-65% in the upper fraction and the non-motile sperm (motility 12.5-32% in the lower fraction (Figure S2a). Standardising OCR values to the number of motile cells shows the differences between “before swim-up” and “upper fractions” are now diminished in two individuals and retained in one (Figure S2b). Therefore, the OCR separation by swim-up is partly but not exclusively related to motility.

Normalizing ECAR in the same way shows a very large range of ECAR per motile sperm (20-160 npH/min/motile sperm) in the Holstein bull before swim-up (Figure S2b, right graph). ECAR is reduced in the upper and lower fraction (40-74 npH/min/motile sperm and 10-32 npH/min/motile sperm, respectively). In the Fleckvieh bull, the upper fraction shows by far higher values than the before swim-up and lower fraction. In the Angus bull, all fractions display relatively low ECAR values. This illustrates that the swim-up does not generally select sperm with higher glycolytic rate (only in Fleckvieh this was the case).

Plotting the average basal OCR over the average basal ECAR (see Figure S1) results in the characterisation of the metabolic phenotype, or energetic potential, i.e. the ability of the cells to change the metabolic pathway and respond to changes in the energy demand. Analysing the metabolic phenotype of the upper and lower fraction (Figure S1) suggests that the upper fraction sperm had more energetic potential, as indicated by the upper fraction moving more to the upper right.

Regardless of the actual values, the ratio of OCR/ECAR can indicate a preference of metabolism for either oxidative phosphorylation or glycolysis^29^. The average OCR/ECAR ratio values of the different fractions are above one, which indicates that bull sperm perform a much higher amount of oxphos compared to glycolysis (Figure S3). The upper fractions show a OCR/ECAR ratio mean value more towards glycolysis than the “before swim-up” and lower fractions (Figure S3) (p<0.0001, see supporting information), indicating a shift in metabolic pathway usage occurs. This difference in OCR/ECAR ratio was consistent across both levels of viscosity (p<0.043, see supporting information). The OCR/ECAR ratio does not represent overall metabolic activity, but rather shows a shift in metabolic pathway usage.

### The influence of medium viscosity

So far, we have analysed metabolic parameters in standard medium. A minimum requirement that the standard medium reflects differences in sperm environments, such as different viscosities, is that swim-up separates fractions that are consistent in their performance across different viscosities. We measured the metabolic rates of the different fractions in two different viscosities (see setup Figure 2a). Adding 1% methyl cellulose to a conventional assay medium results in viscosity values several orders of magnitude higher (at shear rate 5 s^−1^)^24,25^. This viscosity mimics that of fluids of the reproductive tract, reported to be 0.1-1.0 Pa s ^30^ and has previously been used to study sperm motion in viscous environment similar to the *in vivo* conditions^31–34^.

**Figure 2:**
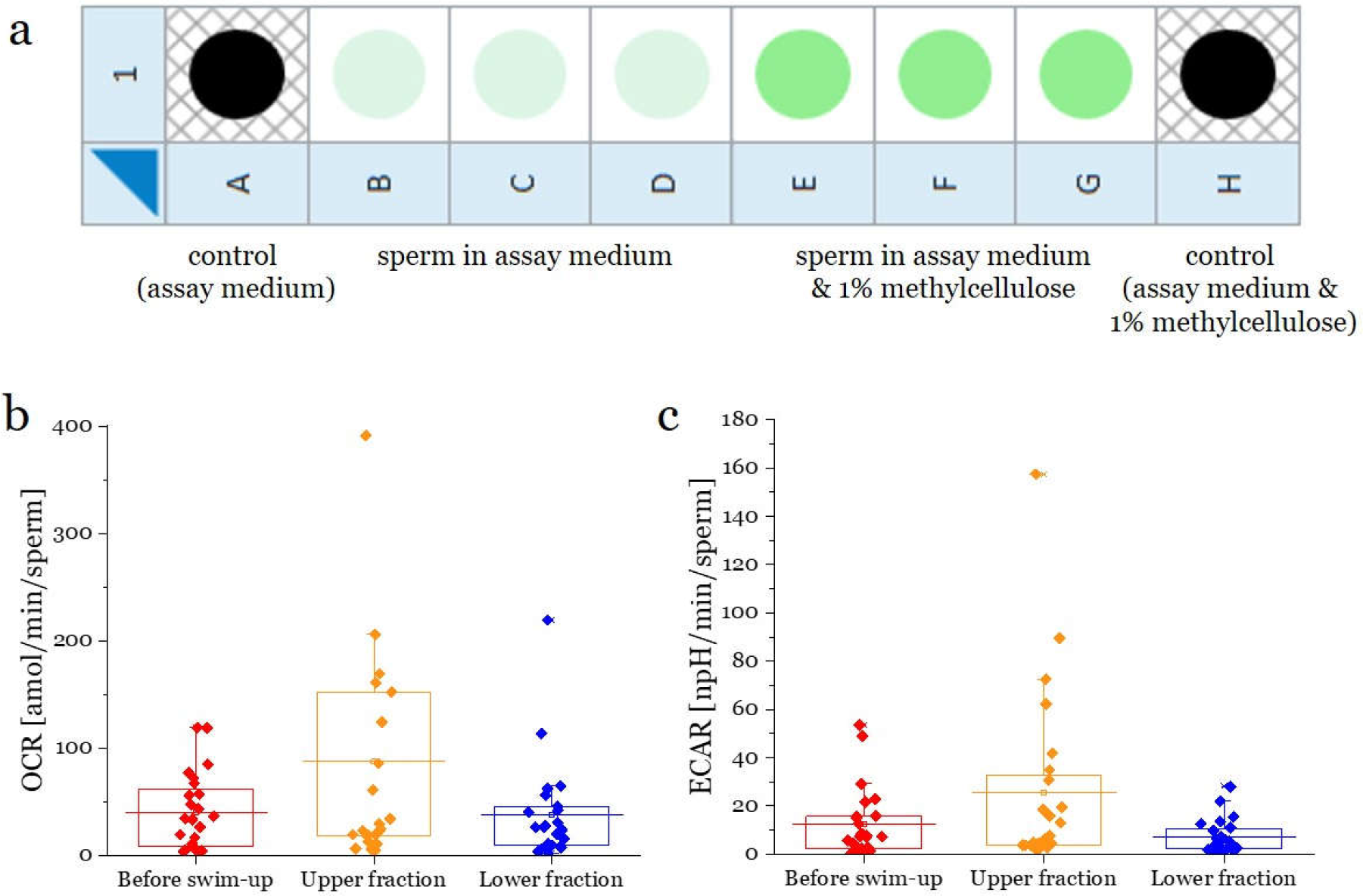
After swim-up in low viscosity medium (sperm medium SP-TALP), oxygen consumption rate (OCR) is compared between low viscosity medium (Assay Medium) and high viscosity medium (Assay medium supplemented with 1% MC). a) Shows the sample distribution in the 8 well plate. This triple-well measurement was performed on each fraction (upper, lower and before swim-up) and two control wells with the two levels of viscosity were used as background level. b) shows the oxygen consumption rate (OCR) and c) shows the extracellular acidification rate (ECAR) for the sperm that did not perform swim-up (before swim-up), the upper and lower swim-up fractions of 12 bulls of different breeds (Holstein, Fleckvieh, Angus). The horizontal lines across the boxes displays the mean values.

Across both viscosities, the lower fractions displayed the lowest OCR (Figure 2b) and sperm with higher OCR accumulated in the upper fractions. Note that the results in standard medium confirm our earlier measurements. There were no differences in sperm OCR rates between low and high viscosity (see Figure S6 in supporting information). This indicates that swim-up selects sperm in the upper fraction that display a higher respiratory activity and therefore have the ability to penetrate fluids of different levels of viscosity. ECAR showed similar values as OCR (Figure 2c), indicating that rates of both pathways are increased in the upper fractions, and therefore, overall metabolic rate.

The difference in energetic phenotype between upper and lower fraction was also maintained through both levels of viscosity (Figure S1), as was the difference in OCR/ECAR ratio (p<0.043, see supporting information Figure S3).

### Swim-up separates sperm into cohorts of high and low ATP production by oxphos

A crucial question is to what degree metabolic differences translate into differences in ATP production. Inhibiting the electron transport chain with Oligomycin A allows a calculation of ATP production rate by oxidative phosphorylation (Figure 3a). Lower fractions (Figure 3b) produced less ATP by oxidative phosphorylation than the upper fractions (p=0.0001, see Table 4 in SI). The upper fractions also had higher values than the initial sperm sample (before swim-up). ATP production was similar in higher compared to lower viscosity for all fractions (Figure S7 in supporting information).

These results indicate that the swim-up separates sperm into fractions of higher and lower ATP production, but that swim-up does not reflect a precise measure of ATP production.

**Figure 3:**
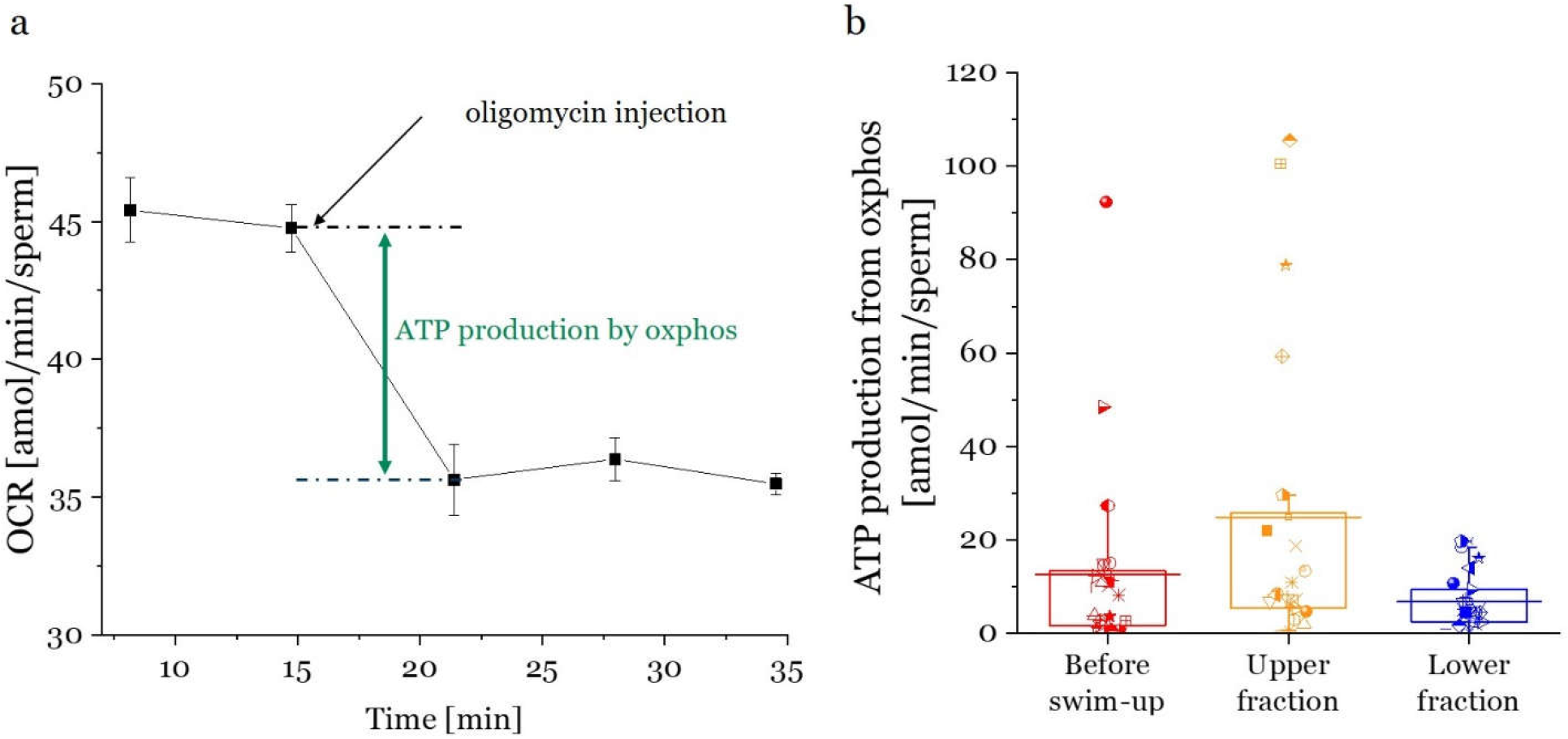
ATP production by oxidative phosphorylation of different swim-up fractions. a) ATP production from oxphos is calculated as the difference between basal OCR rate and OCR rate after oligomycin injection. b) Average ATP production due to oxphos of unselected sperm (before-swim-up), upper and lower swim-up fractions. Each box contains data from 12 bulls. Horizontal lines through each box are the mean values.

Next, we looked at the current ATP content of upper and lower fraction sperm, determined in 11 bulls using a luciferase-based luminescence kit (Figure 4). The bulls showed lower total ATP content in their upper fractions compared to the before swim-up sperm and the lower fraction (p=0.01, see Table 5 in SI). That increased metabolic rates in the upper fraction (Figure 1 and 2) were associated with lower total ATP content (Figure 4) suggests that upper fractions show increased ATP expenditure, perhaps caused by the metabolic activity during swim-up.

**Figure 4:**
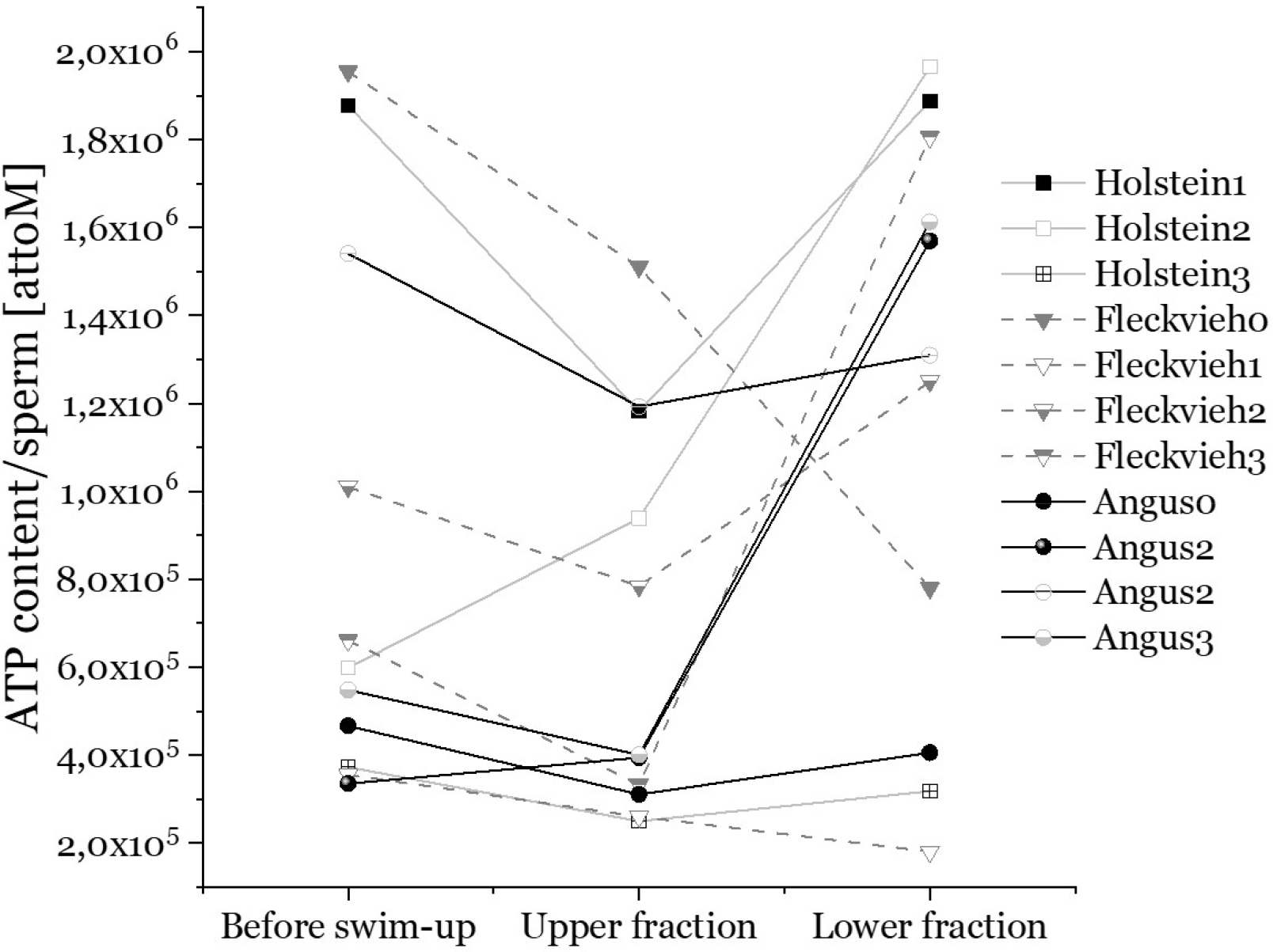
Total ATP content per sperm cell of before swim-up fractions, upper and lower swim-up fractions from 12 bulls measured with a luminescence-based ATP kit. Each condition was measured three times.

Theory suggests that sperm tail length correlates with power output and can thus be considered a selection parameter for most efficient sperm swimming^35^. Indeed, swim-up selected sperm with longer flagella in at least one study^21^. In our study (Figure 5) we found in all individuals that the upper fraction of swim-up sperm contained longer sperm tails compared to the initial sperm sample (p=0.04). The lower fraction consistently contained shorter tailed sperm (p=0.05, Figure 5 and Table 7 in SI).

**Figure 5:**
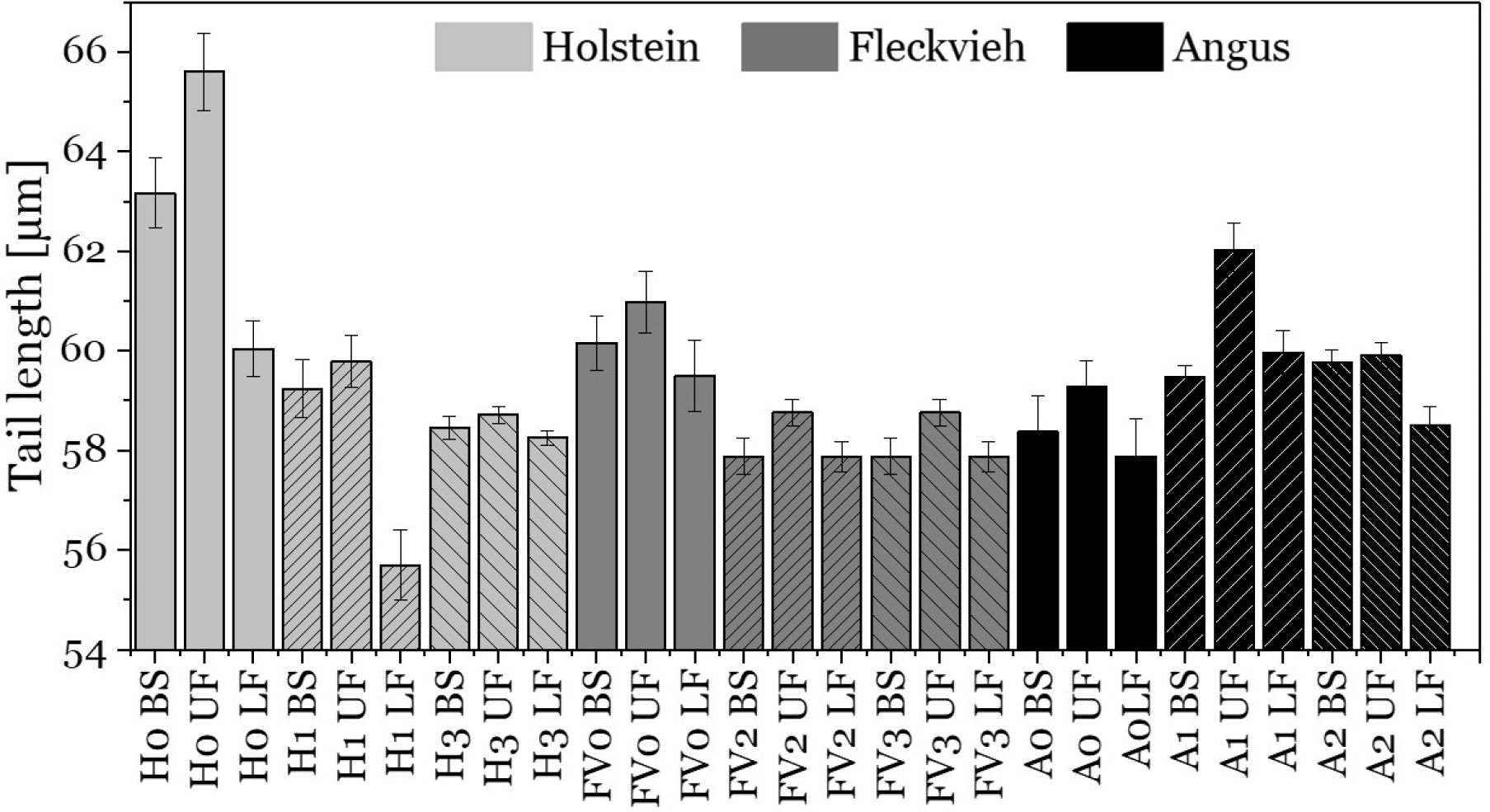
Sperm flagella length of swim-up fractions from 9 bulls of different breeds. 100 sperm cells were measured for each bar. Error bars are standard error of the mean. Upper fractions contain consistently longer sperm tails (estimate= 1.05, p=0.04) compared to the initial sperm sample. The lower fractions contain consistently shorter tails (estimate= −0.98, p=0.05). BS= before swim-up. UF=upper fraction. LF=lower fraction. H0-3= Holstein bulls, FV0-3= Fleckvieh bulls, A0-2= Angus bulls.

## Discussion

This article examines metabolic differences underlying the swim-up method used for sperm analysis in clinical and veterinary settings. We confirmed for bull that swim-up selects sperm of higher motility. We additionally reveal that the standard andrological method of swim-up separates sperm into fractions of high and low metabolic rates and these are based on both, higher oxygen consumption rates and extracellular acidification rates. We also show that swim-up separates sperm of high and low ATP production but that upper-fraction sperm cells have lower ATP content and therefore possibly have higher ATP expenditure. We combined the swim-up method with subsequently immersing the sperm in high viscosity medium and show that the level of viscosity does not significantly change their metabolic rates. Finally, we confirmed a previous notion that because of selecting sperm differing in motility, and as we show in metabolic rate and ATP production, larger sperm are selected.^21^ Overall, this investigation provides a metabolic explanation of why the swim-up method selects sperm that seemingly are functionally superior, suggesting that swim-up is useful in animal reproduction science. It is a possibility that this notion extends to the sperm of other animals and humans.

The upper swim-up fractions displayed both highest oxygen consumption rates and highest extracellular acidification rates. Whether swim-up selects sperm with highest oxygen consumption rates or the highest extracellular acidification rates remains an open question. Our results showing that swim-up selects sperm in the upper fraction with highest ATP production from oxphos may suggest that oxphos plays a key role, but the causal connection between the two remains open.

Our data are consistent with previous studies using different methods in showing a low glycolytic contribution to bull sperm metabolism^5,8^ and we, therefore assume our study to be representative. Whilst the low glycolytic activity argues for a substantial utilisation of oxphos, glycolysis is also discussed as being essential to supply energy to sperm regions far from mitochondria to support motility^7^. For example, inhibiting oxphos while stimulating glycolysis maintained full sperm motility^36^. Other studies suggest that diffusion of ATP in sperm flagella is enough, if there is enough ATP stored in the tail, as it is the case in bull sperm^37^. Additionally, it was observed that sperm cells, especially bovine sperm cells, continue being motile in medium without glucose for a long time, suggesting that gluconeogenesis might play a role in energy supply, yet, clear evidence is lacking.^38^

### Viscosity and its effect on energy expenditure and oxidative damage

Sperm cells in the upper fraction were more likely to penetrate fluids of different levels of viscosity. This supports the hypothesis from simulation-based studies that suggest that sperm cells swim more efficiently in highly viscous media by adjusting their flagellar waveform^39^. It is predicted that, by decreased cell yawing and reduced wavelength, the sperm cell maintains a similar power output in higher viscosity. The change in flagellar waveform induces a greater efficacy (velocity per mechanical power) of the sperm motion, thereby reducing the need for increased energy expenditure. Our study also revealed that increasing the viscosity of the medium does not improve the separation power of sperm cohorts based on metabolic differences, because no differences in metabolic rates were observed in these settings. We, therefore, recommend that higher viscosity should not be used to separate sperm; among others this is to avoid the possibility of the Brynhild effect^40^. This effect applies if separation or quality control thresholds for sperm are set too high, in either the female or in artificial settings, sperm may actually incur reduced functionality or higher energy expenditure and not be useful for fertilisation. For example, a threshold that requires large energy expenditure may select for sperm utilising oxphos, which results in 15 times more ATP compared to glycolysis. However, oxphos, responsible for ~90% of a cell’s ROS production^41^, may increase ROS damage of cell membranes and DNA^41^ thereby reducing sperm function and obsoleting the threshold^42^. Future work should test to what extent upper fraction sperm accumulate oxidative damage.

Longer migration distances of sperm (e.g. from the anterior vagina in case of natural mating) were suggested to be one way of quality control for sperm^43^. While it was argued that such long distances make high metabolic rates beneficial (the winning-the-race argument^43^) it seems that because sperm are stored by females before fertilisation, low metabolic rates may be beneficial because ATP would be used up slower and sperm retain motility for longer. It was predicted that ATP content could be an interesting parameter coupled to fertility and gives supplementary information about sperm quality.^44^ It was reported that ATP content did not necessarily correlate with motility, but was influenced by preservation method and post-thaw incubation time. Yet, further research is needed to understand the connection between ATP content and fertility. In our example, we found that low ATP reserves likely indicated high ATP expenditure. Combining measures of ATP reserves and ATP production rate might be a useful prediction parameter for sperm quality.

In our study we include 12 bulls from three genetically different breeds. The variation in the investigated parameters (OCR, ECAR, ATP content, ATP production, tail length) between bulls was large and therefore prevented an analysis of breed-specific traits. Whilst not the focus of the work presented here, breed-specific sperm metabolism may an interesting aspect of predicting fertility.

Since the swim-up procedure is a common sperm preparation method and a model to mimic sperm migration we think it might be reasonable to test the metabolic differences between upper and lower sperm fractions in human sperm for its potential clinical applicability, especially given that motility and metabolism are not perfectly correlated. An interesting extension would be to assess sperm metabolism in patients with low or no sperm motility or abnormal morphology because these impede the standard motility assays. Further prospects of this method lies in the assessment of metabolism in real time in sperm exposed to drugs, medium additives or toxins.

In conclusion, we propose that measuring metabolic activity of sperm can be an important indicator for sperm quality and their migration success.

## Methods

### Schematic of workflow

Figure 6 illustrates how the separation and subsequent metabolic measurements of bull sperm swim-up fractions were performed.

**Figure 6:**
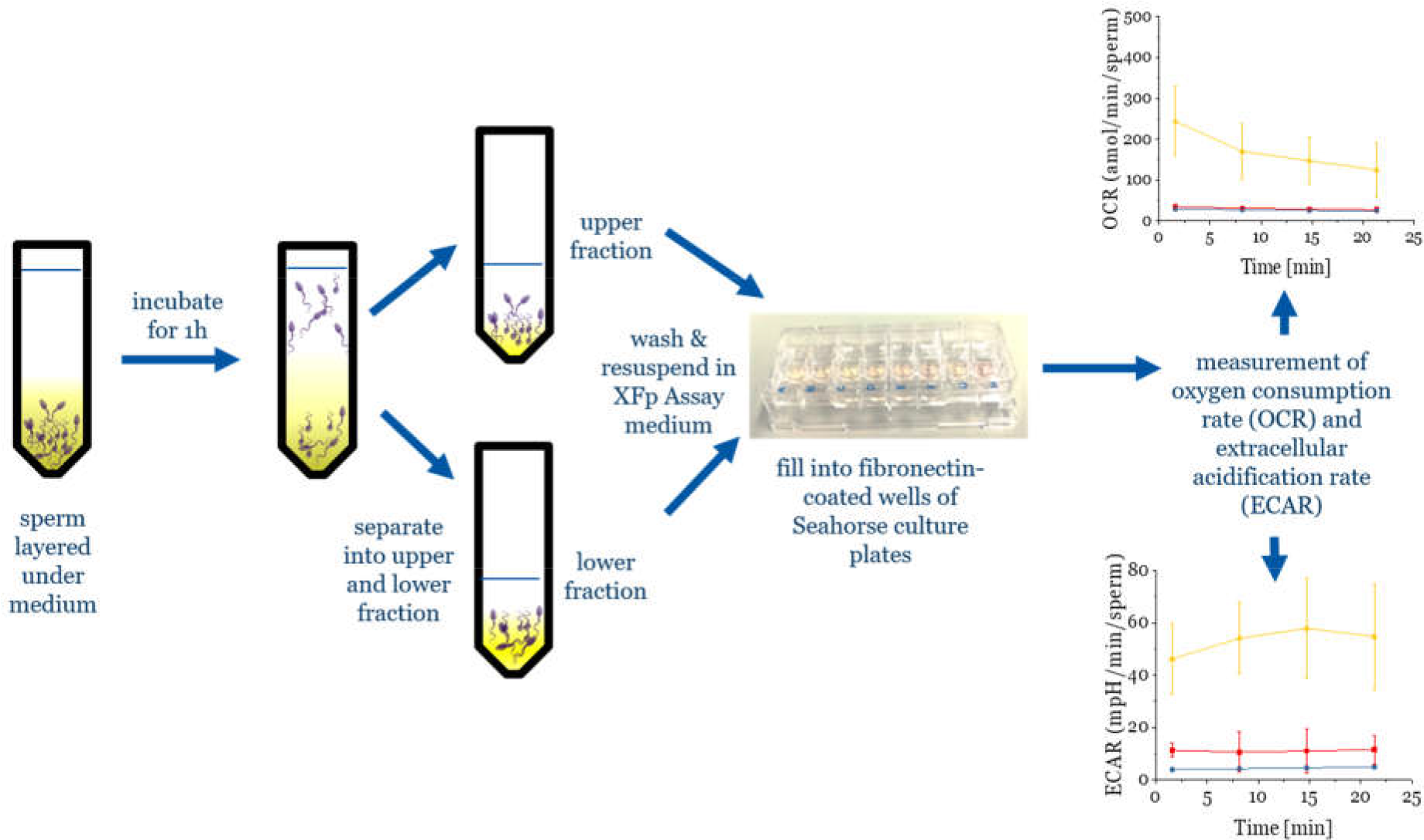
Schematic of the workflow for swim-up of bull sperm and subsequent metabolic measurements.

### Preparation of bovine sperm

Cryopreserved bull semen from 12 bulls from three different breeds (Holstein, Fleckvieh & Angus) was obtained from Masterrind GmbH. The semen straws were thawed in a 38°C water bath for 2 minutes and diluted in 1 mL modified Tyrode’s medium (SP-TALP). SP-TALP was made from SP-TL (Caisson labs), 100mM sodium pyruvate (Gibco), 50 mg/mL Gentamicin (Caisson labs) and 6mg/mL bovine serum albumin (fraction V, Sigma) The vial was centrifuged at 300g for 5 minutes in soft centrifugation mode. The pellet was resuspended in SP-TALP and kept in the incubator (37°C) until further use.

### Swim-up

Swim-up was performed immediately by resuspending the washed sperm pellet in 350µL SP-TALP. The resuspended sperm where then carefully layered under 1 mL SP-TALP in an Eppendorf vial. This vial was placed in the incubator for swim-up for 1 h. Subsequently, the upper 1 mL were withdrawn and placed in a separate vial. Upper and lower fraction of the swim-up were washed once with SP-TALP at 300g for 5 minutes.

### Motility assays

The sperm video recording was performed immediately after swim-up and before the metabolic measurements. Different sperm fractions were placed on glass slide under the microscope (Leica DMI inverted microscope) with heating stage with HT50 control unit (Minitübe) (temperature was set at 38°C). All spermatozoa in observation field (50-300) were recorded for at least 5 sec with a PCC software and Phantom Highspeed camera at 200 fps, using 10x objective magnification and phase contrast condenser. Video records were stored in AVI format prior analyses. Motility analysis was done with the CASA plugin of ImageJ^45^ modified according to Purchase & Earle (2012)^46^ delivering overall motility, average path and straight line velocity (Figure S4).

### Seahorse metabolic assays

Metabolic measurements were performed with a Seahorse XFp device from Agilent with eight well cell culture plates. One day before the assay, the sensors cartridges were hydrated at 37°C in calibration solution (Agilent). On the day of the assay, each well bottom of the cell culture plates was coated with 10 μL fibronectin (1 mg/mL, Sigma F1141) and incubated until dry to allow sperm attachment to the well bottoms (Figure S5). The washed sperm pellet was resuspended in XFp Assay medium (XF Base medium, sodium pyruvate, glutamine, glucose, pH 7.4, sterile filtered) to obtain a cell concentration of 10^5^-10^6^ per well. 50 μL of cell suspension was added to each of six wells (two control wells without cells) and centrifuged for 1 minute at 600 rpm to ensure cell attachment. Assay medium was added to the wells to a total of 180 μL per well. The cell culture plates were kept in the incubator for 10 minutes to acclimatise. Then, the metabolic measurements with Seahorse (Agilent) were started by calibrating the sensor cartridge in the calibration solution. After the calibration, the cell culture plate was inserted. During the Seahorse measurements, the temperature was kept at 37°C and automatic injections were programmed as desired. Oligomycin A injection was performed in order to measure the ATP production due to oxphos. Oligomycin A blocks the proton channel of the ATP synthase, an enzyme necessary for oxidative phosphorylation of ADP to ATP^47^. Thus, the energy production by oxidative phosphorylation is inhibited after injecting Oligomycin A to the bull sperm fractions in the Seahorse device. 20 μL of 10 μM Oligomycin A was injected to each well after basal rate measurement, so that a final concentration of 1 μM Oligomycin A was reached in each well. The ATP production due to oxphos can be calculated by subtracting the OCR rate after injection from the basal OCR rate, as displayed in Figure 3a.

After the metabolic measurements, the viability of cells was checked by viability stain (Live/Dead Sperm viability kit (ThermoFisher L7011, Figure S5 in supporting information) and sperm cells from each well were released by vigorous pipetting and counted in a Neubauer counting chamber in order to normalize the values to sperm numbers.

For the investigation of metabolic rates of bull sperm in two different levels of viscosity, swim-up of bull sperm was performed in sperm medium, as described earlier, then, the different sperm fractions were subsequently immersed in highly viscous medium (by supplementation of 1% methyl cellulose). Methyl cellulose is an inert, biocompatible natural polymer that has been used in various studies to mimic the viscosity of body fluids, also for studying sperm motion^24,25,31,33,34^. Its viscosity is about 100 times higher than conventional sperm medium (at low shear rate of 5 s^−1^), similar to female reproductive tract mucus. Eamer et al. studied sperm motion in dependence of viscosity of medium by preparing different methyl cellulose solutions. Their data show that with increasing viscosity, the average sperm velocity decreases while the sperm linearity increases ^33^. Furthermore, Gonzalez-Abreu et al.^25^ reveal a beneficial effect of raising viscosity of the media on the sperm parameters in terms of higher proportion of fast linear spermatozoa and lower percentage of spermatozoa with slow non-linear movement.

### Luminescence-based ATP kit

Total ATP content was measured with the luminescence-based ATPlite kit (Perkin Elmer). First, an ATP standard curve was set up by preparing an ATP dilution series from 10-^5^ M ATP down to blank. The ATP content of sperm was measured by pipetting 100 μL sperm cell solution into each well of a black 96-well plate, adjusted to 37°C. 50 μL of lysis solution were added and the plate was shaken for 5 minutes. 50 μL substrate solution (luciferase/Luciferin) were added to each well. After shaking for 5 minutes, the plate was dark adapted for 10 minutes and subsequently luminescence was measured.

### Statistical treatment of data

Data were analysed using R version 3.5.2 (2018-12-20)^48^ and the lme4^49^ and glmmADMB package^50,51^. Generalised linear mixed models (GLMM) with negative binomial fits were required because of the skewed data distribution. We tested for the effect of viscosity of the medium (low vs high) and fraction (upper vs lower) on OCR, ECAR and ATP production, and for the effect of fraction on the ATP content of sperm. We used a binomial GLMM to test for the effect of fraction and viscosity on the OCR/ECAR ratio. Initial models contained all possible interaction terms. We reduced the full models stepwise backwards and compared the final models to their respective null models to select the best model. Please see detailed analysis in the supporting information.

## Author contributions

VM, CR & BE conducted experiments on metabolism, motility and tail length. SB analyzed motility data. BE, VM & KR performed statistical analysis. All authors wrote and reviewed the manuscript.

## Acknowledgements

We thank Christin Froschauer and Cornelia Thodte for technical support. We thank the Zukunftskonzept of the TU Dresden, funded under the Excellence Initiative of the German Science Foundation (DFG). SB thanks funding by “CENAKVA” (LM2018099).

