## Supporting Information for "The swim-up technique separates bovine sperm by metabolic rates, motility and tail length"

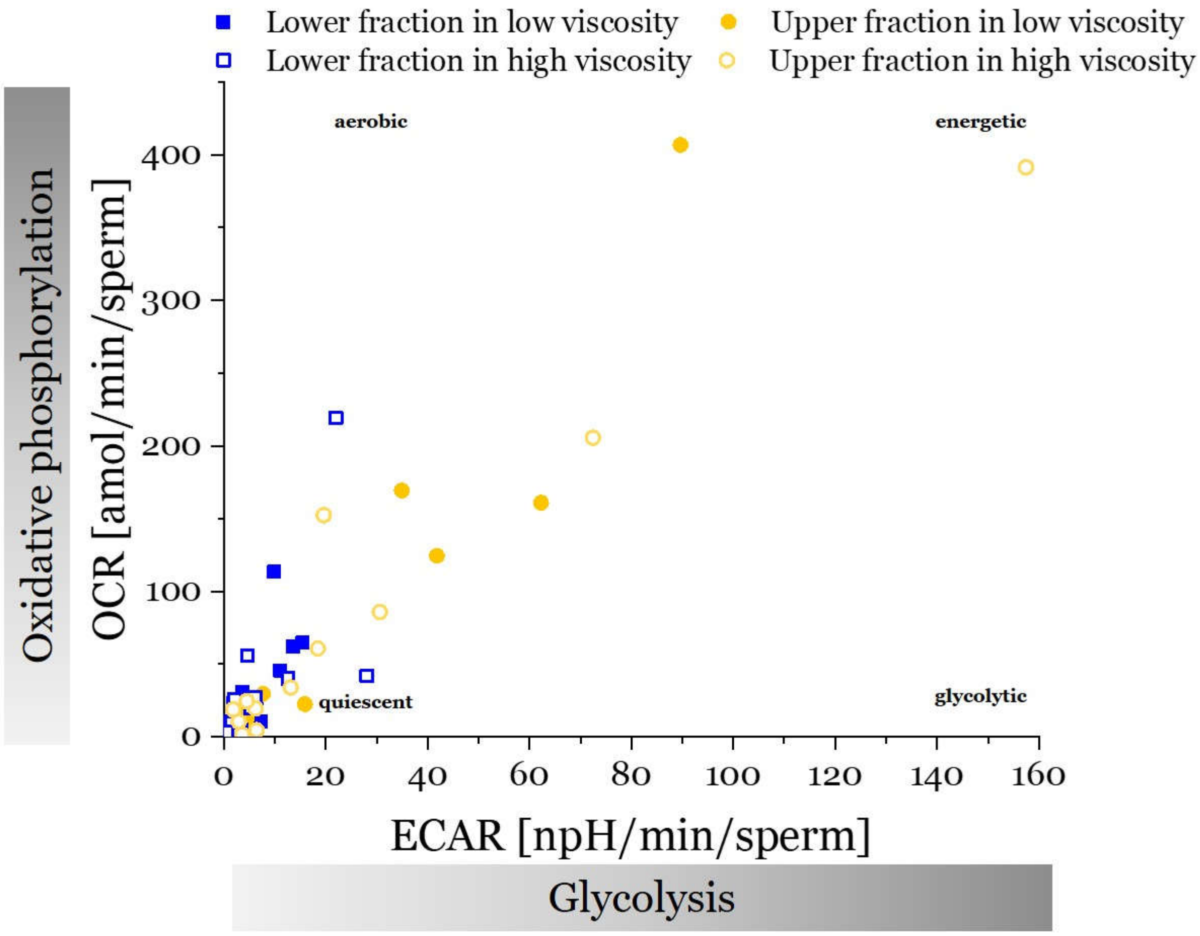


*Figure S1: The metabolic potential is displayed by plotting oxygen consumption rate (OCR) over extracellular acidification rate (ECAR) for the upper and lower swim-up fractions of 12 bulls from 3 different breeds.*

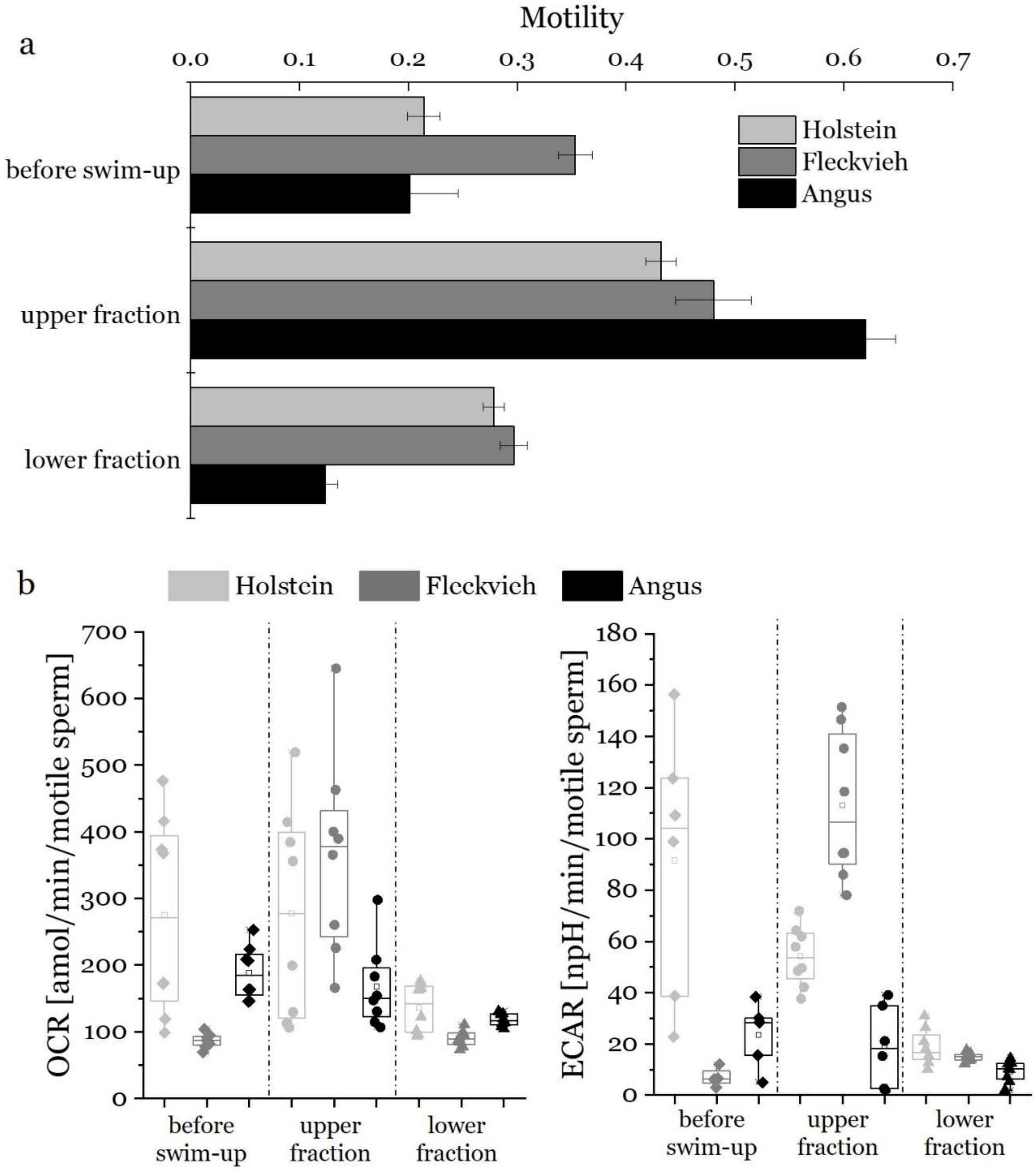


*Figure S2: a) Overall motility of swim-up fractions of bull sperm. b) OCR (left) and ECAR (right) normalized to motility in each fraction. N≥4 replicate measurements.*

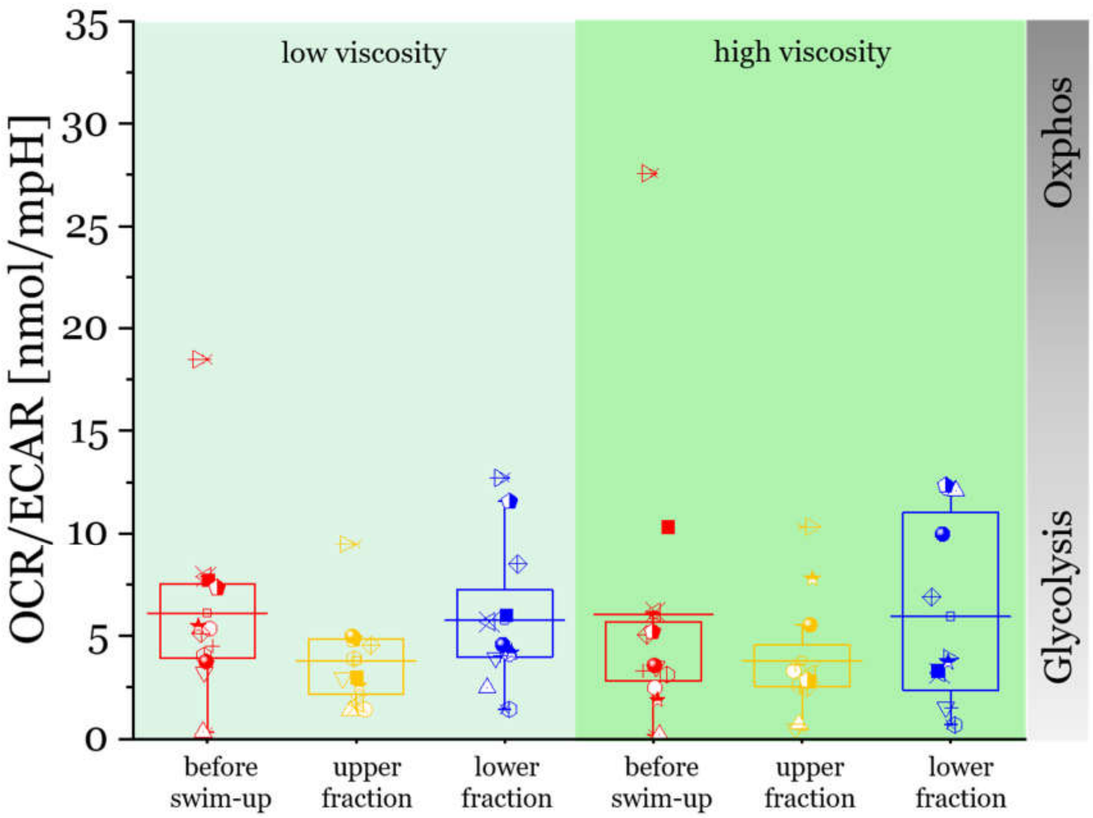


*Figure S3: OCR/ECAR ratios of the sperm fractions before swim-up, upper and lower swim-up fractions. Each box displays OCR/ECAR ratios of 12 bulls, the horizontal line across each box is the mean value.*

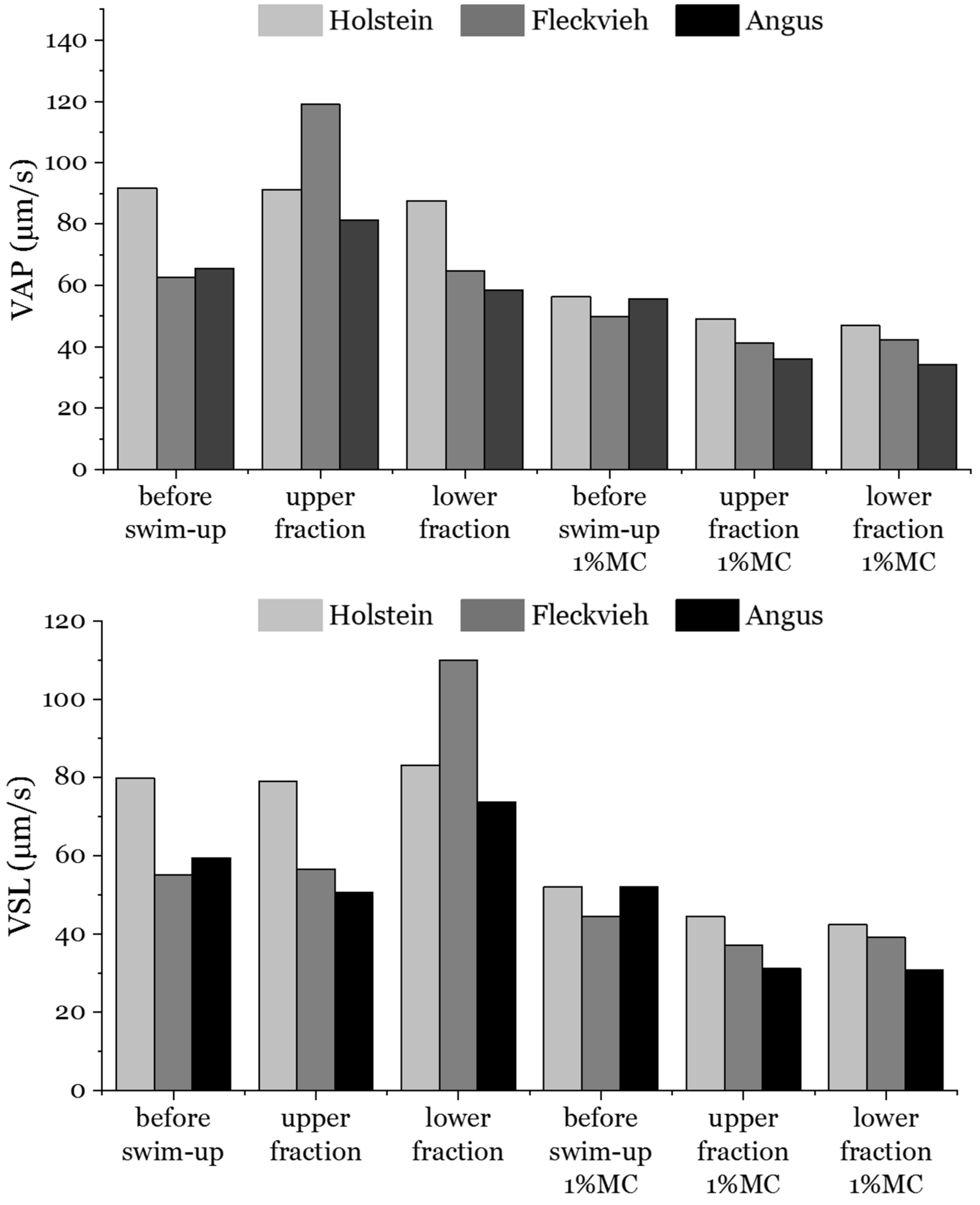


*Figure S4: Average path velocity VAP (top) and VSL (bottom) of different bull sperm swim-up fractions in low viscosity (left) and high viscosity (right) for three different bulls. N >600 sperm cells for each bar. Videos of 10 second length were recorded and then analyzed in one second increments. For each condition and sperm fraction, 3 videos were recorded.*

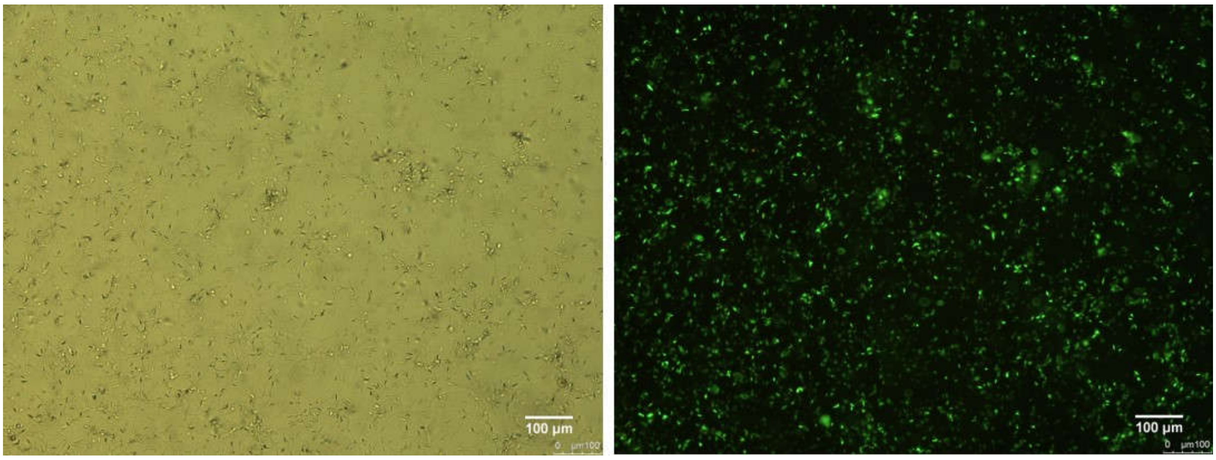


*Figure S5: Sperm cell attachment (left image) during metabolic measurement in the XFp Analyzer and viability stain (right image) after seahorse measurement shows viable sperm cells under assay conditions (37°C, assay medium supplemented with glutamine and glucose).*

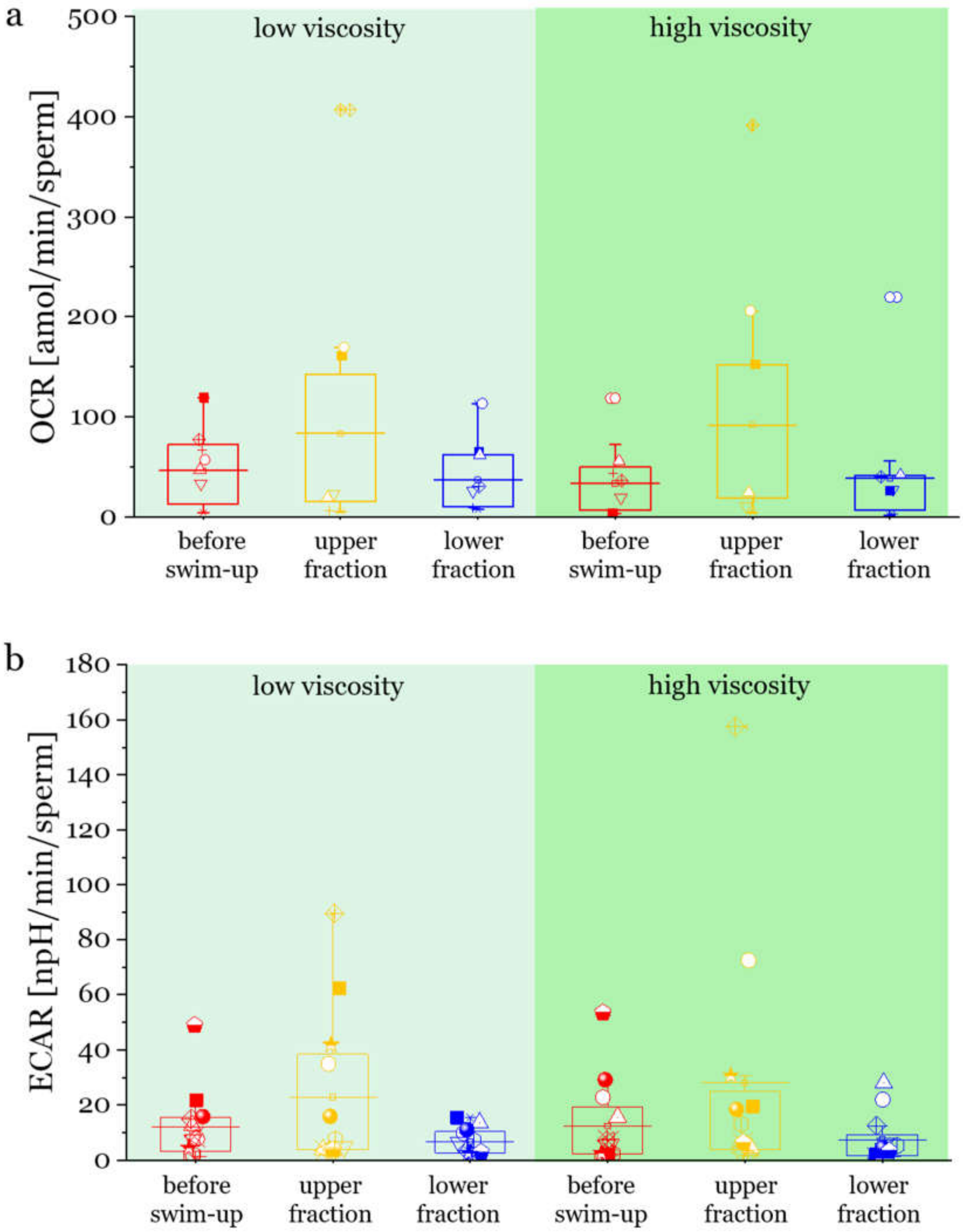


*Figure S6: OCR and ECAR values of the different swim-up fractions across two levels of viscosity. Each box was obtained from 12 semen samples (OCR, p=0.02; ECAR, p=0.00019). Viscosity does not change the OCR or ECAR values significantly.*

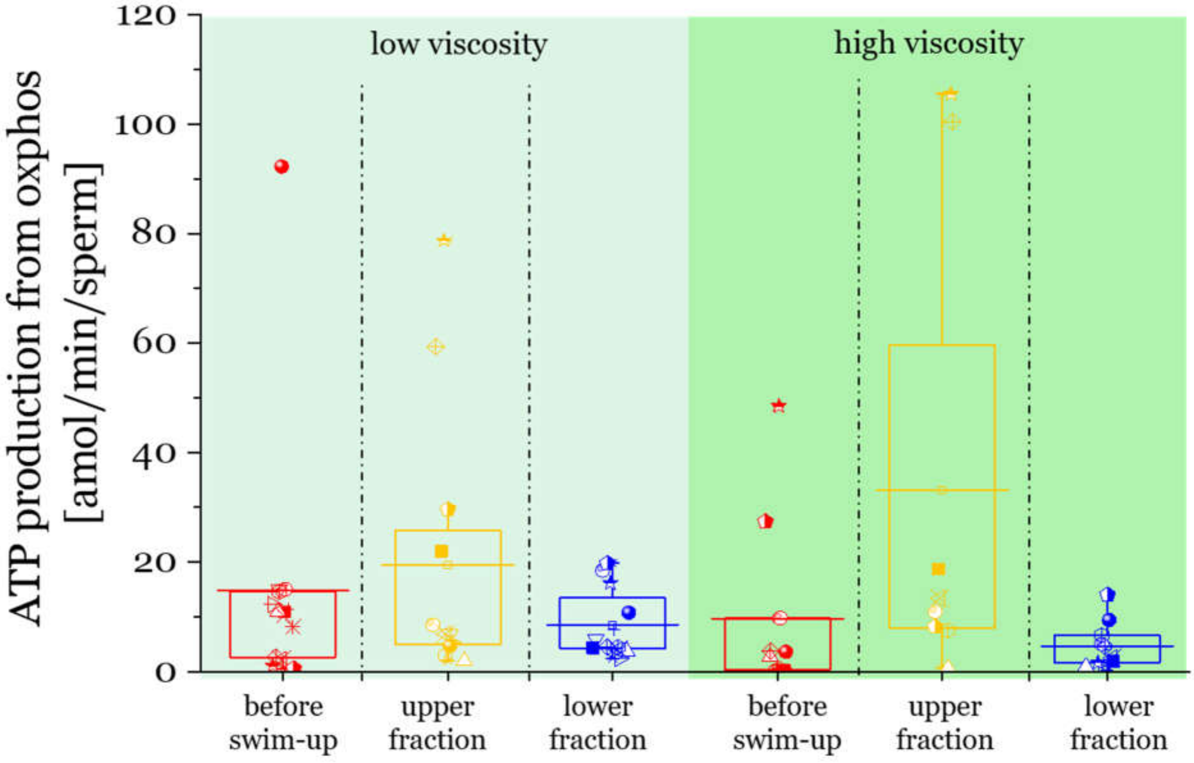


*Figure S7: ATP production from oxphos calculated from oligomycin injections (see Figure 6a) of swim-up fractions in low viscosity (left panel) and high viscosity (right panel). Each box was obtained from 12 semen samples from different bulls. Horizontal lines through each box are the mean values. Viscosity does not influence the ATP production significantly. (p=0.07)).*

### *Results of statistical analysis*

*Table 1 –GLMM (OCR ~ fraction+(1|ID), family="nbinom")*

| **factor** | **estimate** | **SE** | **Z-value** | **p-value** |
|---|---|---|---|---|
| **intercept** | 3.28 | 0.33 | 9.93 | <0.0001 |
| **fraction (upper)** | 0.49 | 0.21 | 2.32 | 0.02 |

*Table 2 –GLMM (ECAR ~ fraction+(1|ID), family="nbinom")*

| **factor** | **estimate** | **SE** | **Z-value** | **p-value** |
|---|---|---|---|---|
| **intercept** | 1.79 | 0.31 | 5.83 | <0.0001 |
| **fraction (upper)** | 0.94 | 0.25 | 3.73 | 0.00019 |

*Table 3 - GLMM (cbind(OCR,ECAR)~fraction+viscosity+(1|ID), family="binomial"l)*

| **factor** | **estimate** | **SE** | **Z-value** | **p-value** |
|---|---|---|---|---|
| **intercept** | 1.60 | 0,12 | 13,60 | <0,0001 |
| **fraction (upper)** | -0.52 | 0,11 | -4,90 | <0,0001 |
| **viscosity (low)** | 0.17 | 0.08 | 2,03 | 0,043 |

*Table 4 - GLMM (ATP production~fraction*viscosity+(1|ID), family="nbinom")*

| factor | estimate | SE | Z-value | p-value |
|---|---|---|---|---|
| **intercept** | 1.39 | 0.36 | 3.92 | <0.0001 |
| **fraction (upper)** | 1.54 | 0.40 | 3.85 | 0.0001 |
| **viscosity (low)** | 0.66 | 0.36 | 1.83 | 0.07 |
| **fraction (upper) * viscosity (low)** | -1.02 | 0.49 | -2.09 | 0.037 |

*Table 5 - GLMM (ATP content~fraction+(1|ID), family="nbinom")*

| factor | estimate | SE | Z-value | p-value |
|---|---|---|---|---|
| **intercept** | 13.87 | 0.22 | 63.77 | <0.0001 |
| **fraction (upper)** | -0.55 | 0.22 | -2.48 | 0.01 |

*Table 6 - LMM (motility~(1|ID))*

| factor | estimate | SE | df | t-value | p-value |
|---|---|---|---|---|---|
| **intercept** | 0.31 | 0.04 | 10.03 | 8.16 | <0.0001 |

*Table 7 - LMM (Tail length~fraction+(1|ID))*

| factor | estimate | SE | df | t-value | p-value |
|---|---|---|---|---|---|
| **intercept** | 59.38 | 0.59 | 12.06 | 100.03 | <0.0001 |
| **fraction (lower)** | -0.98 | 0.46 | 16 | -2.15 | 0.048 |
| **fraction (upper)** | 1.05 | 0.46 | 16 | 2.29 | 0.036 |
